# A bispecific monomeric nanobody induces spike trimer dimers and neutralizes SARS-CoV-2 *in vivo*

**DOI:** 10.1101/2021.03.20.436243

**Authors:** Leo Hanke, Hrishikesh Das, Daniel J Sheward, Laura Perez Vidakovics, Egon Urgard, Ainhoa Moliner-Morro, Changil Kim, Vivien Karl, Alec Pankow, Natalie L Smith, Bartlomiej Porebski, Oscar Fernandez-Capetillo, Erdinc Sezgin, Gabriel K Pedersen, Jonathan M Coquet, B Martin Hällberg, Ben Murrell, Gerald M McInerney

**Author notes:** Contributed equally (Leo Hanke, Hrishikesh Das). Contributed equally (Ben Murrell, Gerald M McInerney).

## Abstract

Antibodies binding to the severe acute respiratory syndrome coronavirus 2 (SARS-CoV-2) spike have therapeutic promise, but emerging variants show the potential for virus escape. This emphasizes the need for therapeutic molecules with distinct and novel neutralization mechanisms. Here we isolated a nanobody that interacts simultaneously with two RBDs from different spike trimers of SARS-CoV-2, rapidly inducing the formation of spike trimer-dimers leading to the loss of their ability to attach to the host cell receptor, ACE2. We show that this nanobody potently neutralizes SARS-CoV-2, including the B.1.351 variant, and cross-neutralizes SARS-CoV. Furthermore, we demonstrate the therapeutic potential of the nanobody against SARS-CoV-2 and the B.1.351 variant in a human ACE2 transgenic mouse model. This naturally elicited bispecific monomeric nanobody establishes a novel strategy for potent inactivation of viral antigens and represents a promising antiviral against emerging SARS-CoV-2 variants.

## Introduction

The SARS-CoV-2 pandemic and the vast societal and economic consequences of the connected lockdowns have had devastating consequences for the world’s populations. Despite rapid progress in vaccine development, some emerging virus variants appear to be less effectively neutralized by antibodies elicited by the first-generation vaccines^1^ and the emergence of further variants is expected. To prepare for different scenarios and to have emergency alternatives at hand, novel ways to inhibit and neutralize the virus and variants of concern are needed. As therapeutic agents in the early phases of SARS-CoV-2 infection, camelid-derived single-domain antibody fragments are promising candidates. These so-called ‘nanobodies’ not only bind their antigen with very high specificity and affinity, but they can also be easily produced to large quantities in *E. coli*^2^ and show favorable biochemical characteristics, including good thermal stability. Importantly, they can easily be combined to form homo- and heterodimers or other multimers to increase potency and target multiple epitopes simultaneously^3–5^. This strategy allows the neutralization of multiple known or unknown virus variants and reduces the risk for viral escape^5^.

The attachment of SARS-CoV-2 to the host cell receptor ACE2 and subsequent membrane fusion is mediated by the viral spike (S) protein^6^. In the prefusion conformation, the spike is a large homotrimer. Each S monomer consists of two subunits: S1 and S2. The S1 subunit contains the receptor binding domain (RBD) and S2 contains the fusion peptide, as well as the transmembrane domain. The RBD is responsible for the initial attachment to ACE2 and is found in two distinct conformational states: The receptor-accessible ‘up’ conformation and the receptor inaccessible ‘down’ conformation^6,7^. Receptor binding induces the rather unstable 3-up state, which leads to shedding of S1 and refolding of S2 thereby presenting the fusion peptide to the opposing membrane^6,8,9^. SARS-CoV-2 is easily neutralized by high-affinity binding antibodies and other spike-targeting molecules, including nanobodies^10–13^. Many of them bind to, or close to, the receptor binding motif of the RBD and neutralize by directly hindering virus attachment^14,15^. Other mechanisms include locking the RBDs in the down conformation or preventing productive fusion^5,16^. Still, the emergence of new virus variants that evade antibody-mediated neutralization mandates the identification of molecules with a more diverse set of binding epitopes and neutralization mechanisms allowing for neutralization of a variety of virus variants. Here, we generated a nanobody that has two non-overlapping interaction sites on the SARS-CoV-2 RBD, each targeting a different epitope, which induces the formation of dimers of spike trimers and rescues mice from lethal SARS-CoV-2 infection.

## Results

### Isolation and characterization of a potent RBD-specific nanobody

To identify nanobodies with unique mode of neutralization, we immunized one alpaca (*Funny*) with recombinant SARS-CoV-2 spike protein and RBD. We isolated RNA from peripheral blood mononuclear cells and created phagemid libraries. Phage display screens with immobilized RBD identified one promising nanobody, called Fu2. Since multimerization of nanobodies can substantially improve virus neutralization potency^3,17^, we created three different dimers (illustrated in Fig. 1A): An Fu2-Fc fusion expressed in mammalian cells, a chemically linked Fu2 homodimer, and thirdly, a heterodimer of Fu2 with a previously described RBD-specific neutralizing nanobody Ty1^14^. Chemically-linked constructs were generated using a combination of sortase A labelling and Cu-free click-chemistry as described in detail previously^3^.

**Fig. 1.**
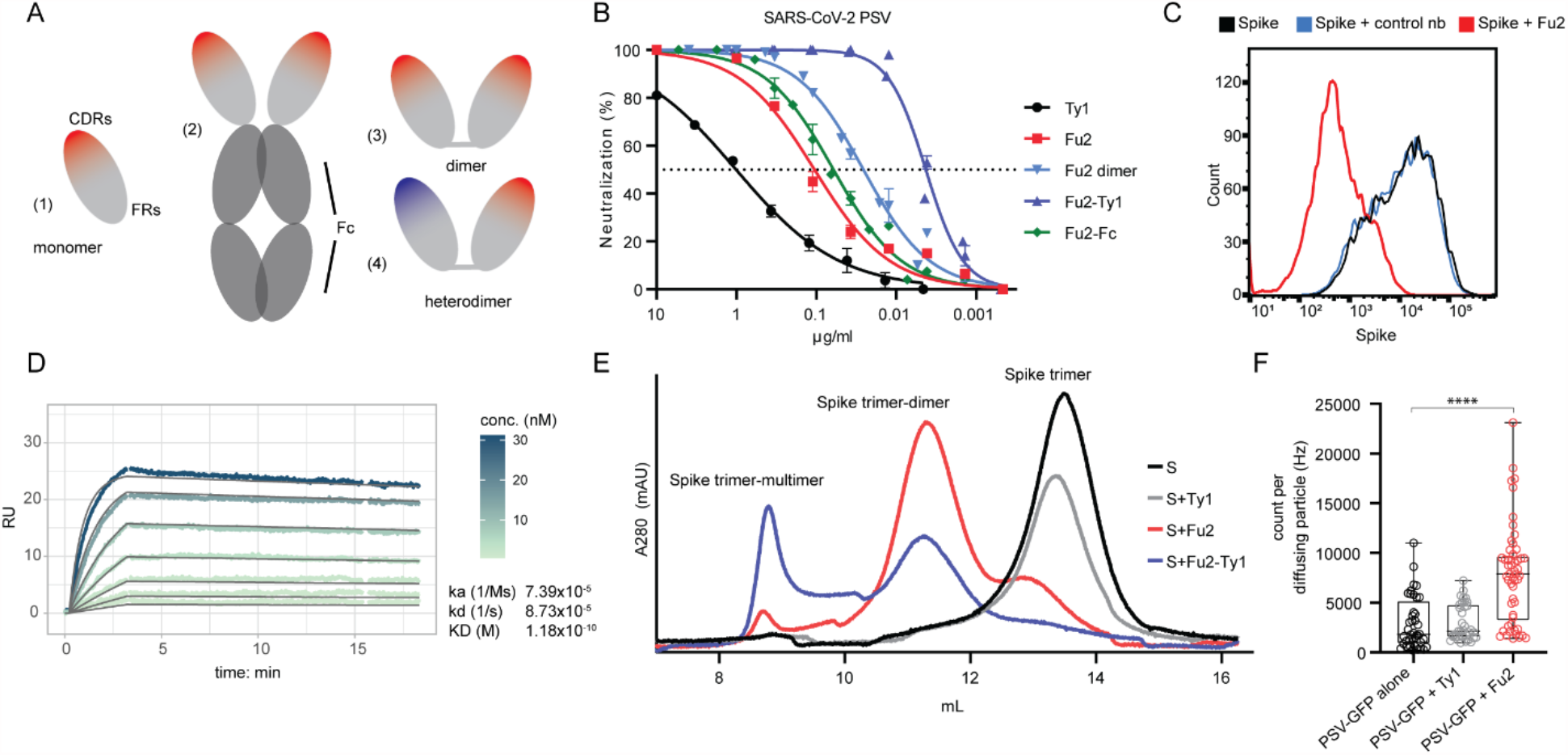
An RBD-specific nanobody neutralizes SARS-CoV-2. (**A**) Overview of different nanobody constructs used in this study: (1) nanobody monomer, (2) nanobody Fc-fusion, (3) chemically linked nanobody homodimer, (4) chemically linked nanobody heterodimer. (**B**) A SARS-CoV-2 spike pseudotyped lentivirus (PSV) was incubated with a dilution series of the indicated nanobodies at 37°C for one hour before infecting HEK293T-hACE2 cells. Neutralization (in %) compared to untreated PSV is shown. Data from at least two replicates is shown and the error bars represent the standard deviation. (**C**) Recombinantly expressed, prefusion stabilized, and fluorescently labelled SARS-CoV-2 spike protein was incubated with a control nanobody specific for IAV NP^31^, or Fu2 and used to stain ACE2 expressing HEK293T cells. Cells were analyzed by flow cytometry and a representative histogram is shown. (**D**) Binding kinetics of Fu2 to the RBD were measured by surface plasmon resonance (SPR). Sensorgram is color-coded based on concentration. The fit is based on the 1:1 Langmuir model and is shown in dark grey solid lines. (**E**) Recombinantly expressed, prefusion stabilized SARS-CoV-2 spike protein was run alone or preincubated with the indicated nanobody constructs on a Superose 6 size-exclusion column. Elution profiles (A280) are shown. (**F**) GFP-labelled SARS-CoV-2 spike pseudotyped lentivirus (PSV-GFP) was incubated with Ty1, Fu2 or left alone and brightness of diffusing virus particles was quantified using fluorescence correlation spectroscopy in solution. Statistical significance was evaluated using Kolmogorov-Smirnov test.

Using a pseudotyped lentivirus (PSV) neutralization assay (Fig. 1B), we found that monomeric Fu2 was approximately 10-times more potent than Ty1, with an IC_50_ in the range of 106 ng/ml (7 nM). The Fu2-Fc fusion only slightly increased the neutralization potency of this nanobody to 61 ng/ml (0.75 nM), and a chemical homo-dimerization to an IC_50_ of 26 ng/ml (0.8 nM). Thus, the observed enhancement in potency is attributable to dimerization, rather than the Fc-domain itself. Interestingly, the Fu2-Ty1 heterodimer was extremely potent, with an IC_50_ of 4 ng/ml (140 pM) which is approximately 6-fold lower than that of the Fu2 homodimer. Fu2 directly prevented spike binding to ACE2 as confirmed by flow cytometry of HEK293T-hACE2 cells stained with recombinant, fluorescently labelled spike (+/- Fu2) (Fig. 1C). Analysis of binding kinetics by surface plasmon resonance (SPR) revealed that Fu2 bound the RBD with sub-nanomolar affinity (Fig. 1D). A 1:1 binding model fit the sensorgrams well at the used concentration range, indicating a single dominant interaction. Analysis of next-generation sequencing data for other members of the Fu2 lineage yielded several Fu2-variants with similar properties (Figs. S1A-E).

To test the integrity of the recombinant spike protein after the addition of Fu2, we analyzed the size-exclusion chromatography elution profiles in combination with different nanobody constructs (Fig. 1E). The SARS-CoV-2 spike trimer eluted in a single peak and the addition of Ty1^14^ did not result in a shift in the position of this peak, which is expected given the small relative size addition of the 14 kDa nanobody to the large (>400 kDa) spike trimer. In contrast, the addition of Fu2 resulted in a substantial shift of the peak, suggesting the formation of a larger complex, likely a dimer of the spike trimer. Next, we combined the spike with the Fu2-Ty1 heterodimer. This also resulted in earlier elution, and besides a possible spike dimer, this construct gave rise to an even earlier elution peak suggesting the induction of higher-order spike multimers.

To test if the Fu2-induced multimerization of spike also occurs when it is presented on virus particles, we analyzed GFP-labeled SARS-CoV-2 pseudotyped lentiviruses (PSV-GFP) after addition of the nanobodies Ty1 or Fu2 (Fig. 1F). To this end, we performed fluorescence correlation spectroscopy and measured apparent brightness of each diffusing particle. The majority of single diffusing particles preincubated with Fu2 showed a 2.5-fold increase in brightness compared to PSV-GFP alone or preincubated with Ty1, suggesting that Fu2 leads to aggregation of the PSV-GFP particles.

### Structural basis for the induction of spike trimer dimers by Fu2

To better understand neutralization and the aggregation of spike by Fu2, we used electron cryomicroscopy (cryo-EM) to determine the structure of the Fu2-spike complex. The structure revealed a remarkable head-to-head dimerization of the trimeric spike, bound by six molecules of Fu2 (Fig. 2). This distinct structural state appeared as prominent 2D classes in our analyses (Fig. 2A). Binding of Fu2 to the RBD is unique as it can simultaneously interact with two different RBDs from different spike trimers in the up conformation (Figs. 2B and 2C). To our knowledge, Fu2 is the only nanobody or spike-binding molecule developed against the SARS-CoV-2 spike to have this property. Our high resolution (2.9Å; 0.143 FSC) localized reconstruction of the RBD-Fu2-RBD-Fu2 interface helped us elucidate the molecular details of this interaction. Each RBD in the complex provides two binding interfaces for two Fu2 molecules, which we call ‘interface-major’ and ‘interface-minor’ according to surface area covered (Figs. 2D, S2A and S2B). The interface-major and interface-minor have a buried surface area of ~740 Å^2^ and ~495 Å^2^, respectively. Parts of the Fu2 framework regions drive the dimerization by shape-complementarity and coupled solvent exclusion in the two interfaces. In addition, the interface-major consists of 15 hydrogen bonds and one salt bridge between Fu2 D117 and RBD K378, while the interface-minor comprises five hydrogen bonds. Furthermore, a region around the Fu2 β-turn A40-K43 plays a key role in dimerization through interactions with two proximal RBDs (Figs. S2C and S2D). Specifically, one of the two proximal RBDs’ T500-Y505 region packs against the Fu2 A40-K43 β-turn and the RBD Y505 is in addition bound by Fu2 Q39. Meanwhile, the other proximal RBD is bound through a bidentate hydrogen bond between Fu2 E44 and the backbone amides of RBD V503-G504 (Figs. 2E, S2C and S2D). Interestingly, the overall structure of Fu2-bound spike had no significant deviations from previous structures (Fig. S3), and we did not find any contact between any two Fu2 molecules or between spike trimers suggesting that the observed spike trimer dimerization is solely driven RBD-Fu2 interactions.

**Fig. 2.**
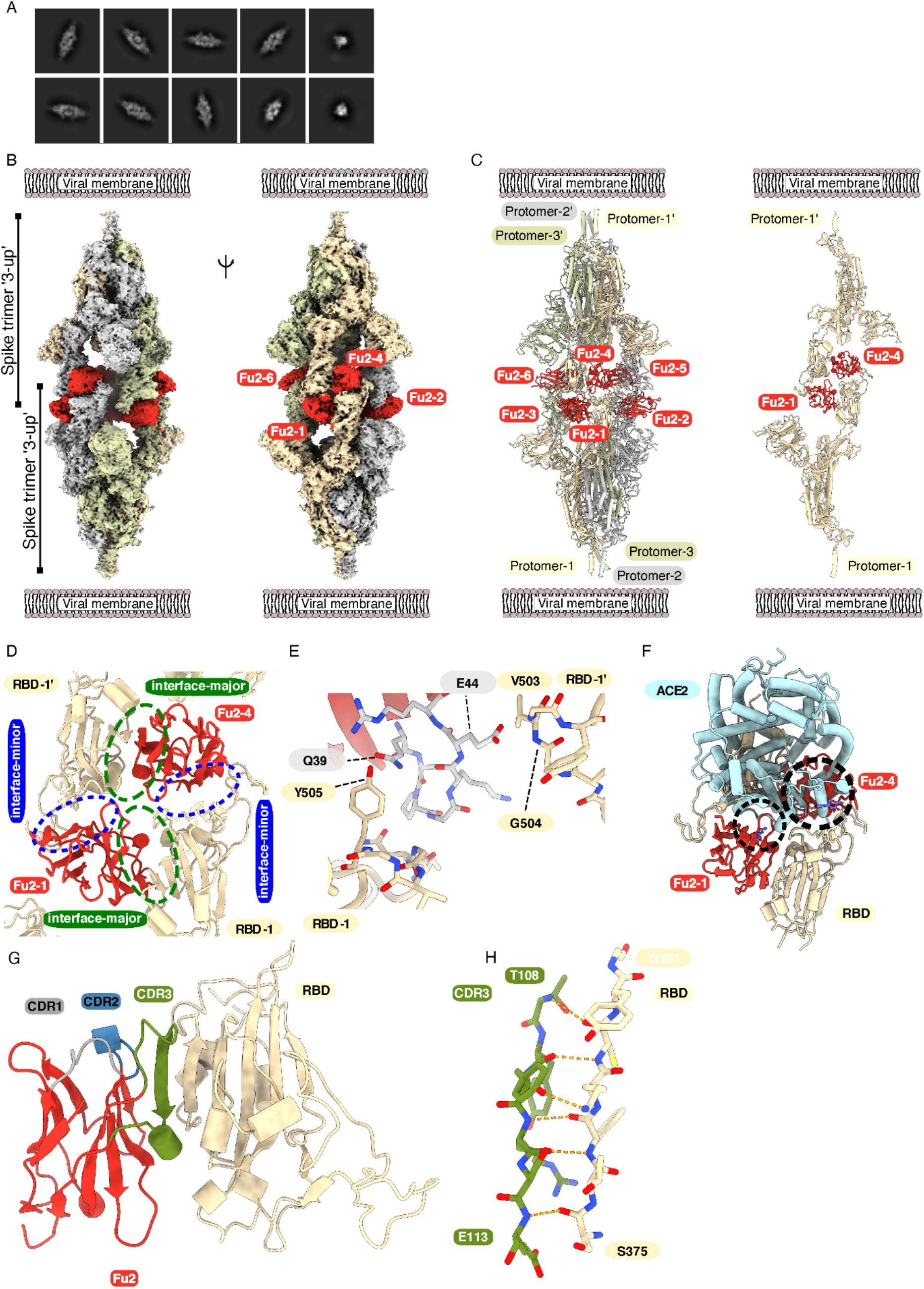
Cryo-EM provides structural evidence of Fu2-mediated dimerization of trimeric spike in ‘3-up’ conformation. (**A**) Representative 2D class averages. (**B**) Cryo-EM map (two different side views) and (**C**) Atomic model of ‘dimeric’ spike in complex with Fu2; and two opposite protomers with two Fu2 molecules. Both spike trimers in the complex are present in ‘3-up’ conformation (six Fu2 molecules are numbered). (**D**) Close-up view of the RBD-Fu2 interaction interfaces showing the interface-major and interface-minor. (**E**) A β-hairpin (gray sticks) from the Fu2 framework region (residues 39-45) contacts two RBDs simultaneously in the Fu2-mediated spike trimer-dimer. (**F**) Modelling of simultaneous spike-Fu2-ACE2 binding shows that Fu2 blocks the binding of ACE2 (PDB:6LZG)^18^ to the RBD from two directions. Clashes marked with dashed circles. (**G**) Fu2 binds the RBD through an extended inter-chain β-sheet interaction mediated by the CDR3 region (green). CDR1 (gray) and CDR2 (blue) does not take part in RBD binding (**H**) Closeup of a part of the CDR3-RBD interface displaying the anti-parallel β-strand hydrogen bond pairing between the Fu2 CDR3 region and the RBD (light orange).

### Fu2 blocks the RBD from binding ACE2

Structural alignment of an ACE2-RBD structure (PDB: 6LZG^18^) to the Fu2-spike structure (single RBD) shows that binding of ACE2 would be hindered by Fu2 bound to any of the two interfaces (Fig. 2F). Fu2 bound to the interface-major would clash with ACE2 residues ~303-331 and the N-glycan at position 322, while Fu2 bound to interface-minor would mask the interaction surface of ACE2, making it sterically impossible for ACE2 to access the RBD. A comparison of the Fu2 and Ty1 binding site revealed that interface-minor is partially shared between Ty1 and Fu2 (Figs. S2E and S2F), suggesting that in context of the Fu2-Ty1 heterodimer, Ty1 is displaced upon dimerization of the spike trimer. Ty1 can then potentially bind additional spike molecules, explaining the elution profile showing both dimers and higher order spike complexes (Fig. 1E).

In the complex, Fu2’s long CDR3 (G99 to Y120) plays a significant role in RBD binding. Here, the Fu2 S107-R111 region adopts a β-strand conformation that binds the RBD T376-Y380 β-strand to give rise to an energetically favorable inter-chain extension of the RBD anti-parallel β-sheet (Figs. 2G and 2H). Interestingly, the Fu2 CDR1 (G26-Y32) and CDR2 (T52-S57) do not participate in RBD binding, and none of the CDRs is directly involved in the formation of the spike dimer.

### Fu2 stabilizes the RBD in the ‘up’ conformation

Cryo-EM analyses, both of the purified spike or of intact virions, have revealed that unliganded spike is mostly present in all-down (~60%) or 1-up (~40%) conformation^6^. Our Fu2-spike structure shows the presence of two trimeric spike molecules, both in the 3-up conformation. This arrangement allows Fu2 to access all six possible binding sites on RBDs without steric conflict. Alignment of RBD-Fu2 to RBD-down in a 2-up spike structure confirms that binding of Fu2 to RBD in the down conformation is not possible (Fig. S2G). Fu2 bound to either interaction interface on an RBD in the down conformation would clash with neighboring RBDs. Likewise, an alignment of the 2-up spike structure to the Fu2-trimer-dimer shows that RBD-down would clash with the neighboring Fu2 molecule (Fig. S2H). In conclusion, for Fu2 binding and for trimer-dimer assembly, all RBDs must be in the up conformation.

### Fu2 cross-neutralizes SARS-CoV, and clinical isolates of SARS-CoV-2 including the B.1.351 variant of concern

To test the breadth of neutralization, we evaluated cross-neutralization of SARS-CoV PSV (Fig. 3A). While Ty1 did not neutralize SARS-CoV, Fu2 and dimeric Fu2 constructs were neutralizing, but less so than against SARS-CoV-2 PSV. Interestingly, the neutralization curves displayed a much flatter slope as compared to those against SARS-CoV-2 PSV. It required 6.1 μg /ml of monomeric Fu2 to reach 50% neutralization in this assay, while 570 ng /ml of Fu2-Fc and 57 ng/ml of the Fu2-dimer was sufficient for 50% neutralization.

**Fig. 3.**
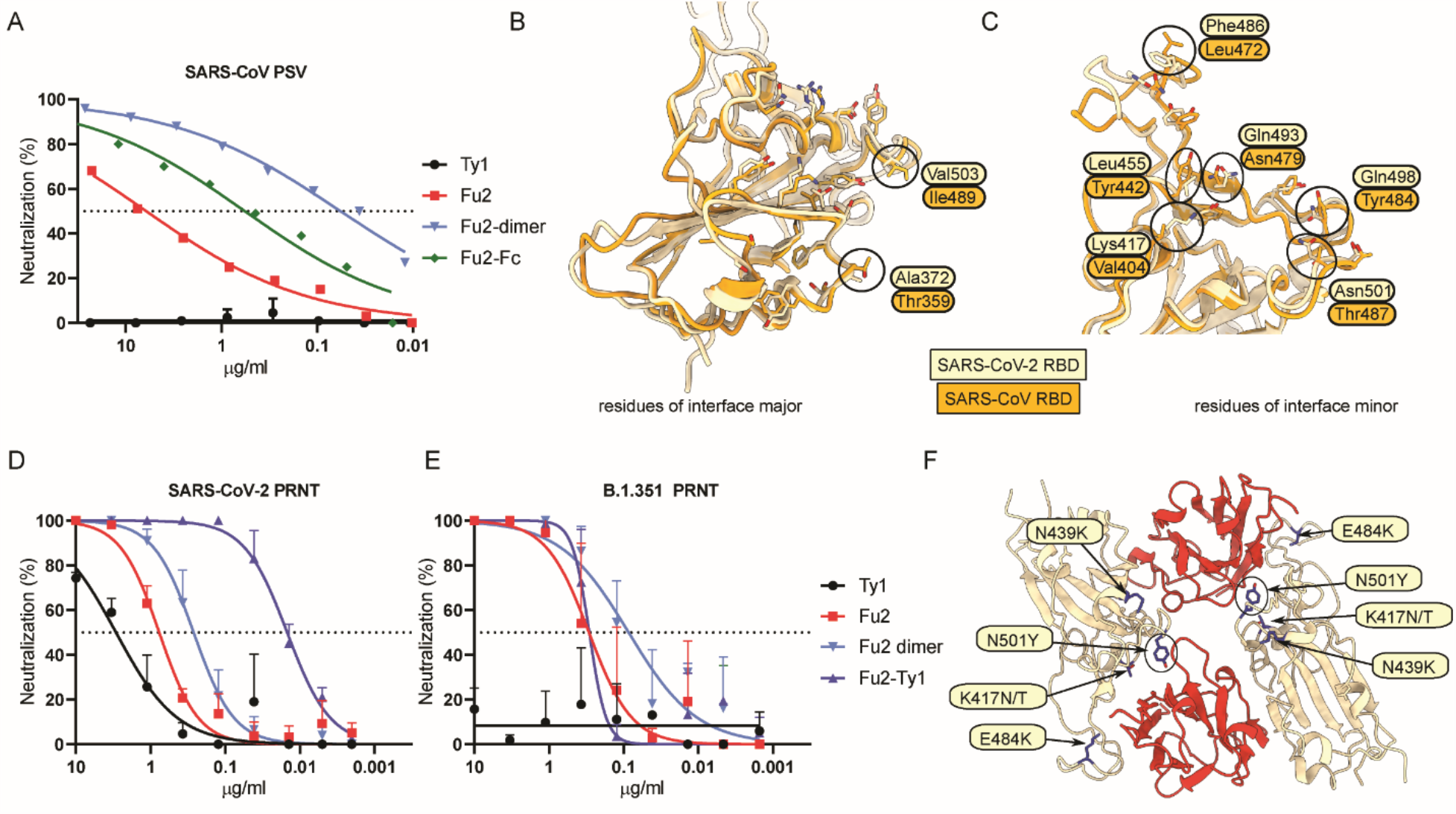
Fu2 cross-neutralizes SARS-CoV and a B.1.351 variant of concern clinical isolate. (**A**) SARS-CoV S pseudotyped lentivirus (PSV) was incubated with the indicated constructs before infecting HEK293T-ACE2 cells, and neutralization is shown compared to untreated PSV. (**B** and **C**) Structural basis of SARS-CoV cross-neutralization by Fu2. Structural alignment of SARS-CoV-2 Spike (RBD bound Fu2) with SARS-CoV RBD (PDB:6ACD)^32^. Residues of interface-major and interface-minor shown as sticks, divergent residues are labelled. (**D and E**) PRNTs with clinical isolates of SARS-CoV-2. 100 plaque forming units (PFU) of SARS-CoV-2^19^ or B.1.351 (501Y.V2)^1^ were incubated with dilution series of the indicated nanobody constructs for 1 h at 37°C before infecting monolayers of Vero E6 cells. Plaques were quantified 72 h post infection. Neutralization representing the reduction of plaques relative to the control wells is shown. Error bars represent standard deviation (SD) across replicate experiments. (**F**) Assessment of Fu2-RBD interaction in RBD variants. *In silico* mutational analyses of RBD residues and probable impact on Fu2 binding. Fu2 is shown in red and variant residues are indicated.

To understand the molecular mechanism of cross-neutralization of SARS-CoV we compared the RBD sequences and performed structural alignments (Figs. 3B, 3C and S4A). Interestingly, the interface-major residues are conserved, while interface-minor has significant differences, suggesting that the divergent interface-minor of SARS-CoV RBD would not permit Fu2 binding. From these analyses we conclude that Fu2 would be unlikely to induce SARS-CoV trimer dimers.

To demonstrate neutralization of replication competent SARS-CoV-2 we performed a plaque reduction neutralization test (PRNT) using infectious SARS-CoV-2, isolated from the first Swedish COVID-19 patient^19^ (Fig. 3D). Similar to neutralization of PSV, monomeric Fu2 neutralized SARS-CoV-2 much better than Ty1, a dimeric construct of Fu2 provided a 2.8-fold improvement over the monomeric version of Fu2, while the heterodimer of Ty1 and Fu2 showed a 50-fold improvement compared to monomeric Fu2 in this assay.

A PRNT using the B.1.351 (501Y.V2) variant^1^ showed that Fu2 very potently neutralized this variant (Fig. 3E) and that the Fu2 dimer and Fu2-Fc again improved neutralization efficiency. The IC_50_ values were even slightly better than when using the Swedish isolate, but the difference is within the intraassay variability. Ty1 failed to neutralize this virus, and the Fu2-Ty1 heterodimer did not neutralize better than Fu2 alone. The variant B.1.351 harbors four amino acid substitutions in the RBD at position K417N, N439K, E484K and N501Y. Only N501 and E484 are in proximity of the Fu2 interaction interface. *In silico* analyses with the PISA server (25) confirmed that the two interfaces do not change in these mutants. Both buried surface areas of the interfaces as well as the number of possible hydrogen bonds remain unchanged (Fig. 3F). These results and analysis combined suggests that Fu2 likely also neutralizes the recently emerged variants P1 (K417T, E484K, N501Y) and B.1.1.7 (N501Y).

### Therapeutic efficacy of Fu2 in a mouse model of SARS-CoV-2 infection

An important requirement for the development of antivirals is neutralization potential *in vivo*. To address this, we used K18-hACE2 transgenic mice that express human ACE2 as a model for SARS-CoV-2 infection. To determine if Fu2 could reduce disease severity when administered in early infection, we tested its ability to neutralize the virus and protect mice from signs of disease *in vivo* (Fig. 4). When the Fu2-Ty1 heterodimer was administered prophylactically or therapeutically, we noted that it slightly delayed the onset of disease (Fig. S7). A major issue with using nanobodies therapeutically is their short half-life *in vivo*. To extend serum half-life, we fused Fu2 to the nanobody Alb1 that binds mouse serum albumin^20^. Mice were challenged with a standardized dose (2.4×10^6^ RNA copies) of either a Swedish isolate of SARS-CoV-2 (86 plaque forming units, PFU) or the B.1.351 variant (100 PFU). On days 1, 3, 5 and 6 post infection, mice were injected intraperitoneally (i.p.) with 600 µg of the Fu2-Alb1 heterodimer (Fig 4A). All control untreated mice experienced considerable weight loss within the first week of infection, with mice infected with B.1.351 experiencing a slightly faster onset of weight loss. In contrast, Fu2-Alb1 delayed disease onset and protected mice from severe weight loss following infection with either the wild-type or B.1.351 virus (Fig. 4B and 4C). Quantification of genomic and subgenomic SARS-CoV-2 E-gene at day 5 also demonstrated that Fu2-Alb1 reduced viral loads, most clearly in mice infected with wild-type virus (Fig. 4D). Thus, Fu2 is a suitable candidate for antiviral therapy against SARS-CoV-2 and common variants.

**Fig. 4.**
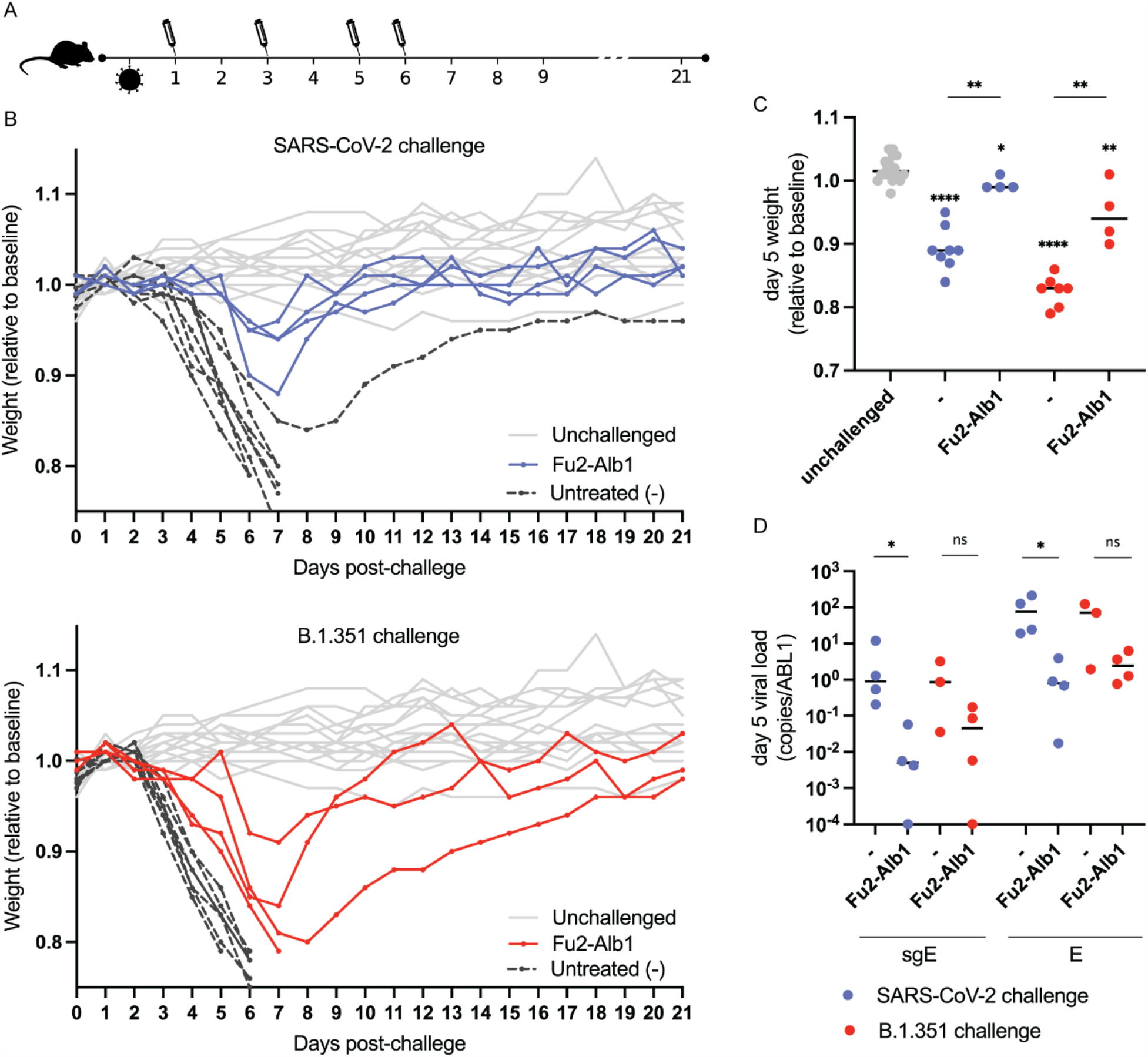
Therapeutically administered nanobody rescues mice from lethal SARS-CoV-2 infection. (**A**) Timeline of the challenge experiment. K18-hACE2 transgenic mice were challenged with 86 PFU of SARS-CoV-2 (blue, ‘wild-type’) or 100 PFU of B.1.351 (red) and injected with Fu2-Alb1 (half-life extended Fu2) at indicated time points. (**B**) Weight of mice during the challenge experiment. The mean weight of each mouse of day 0 to day 2 served as baseline and the weight loss relative this baseline is shown. Uninfected mice are shown in grey, untreated infected mice in black. Half of the untreated mice are historical controls from identical, previously conducted experiments. The weight for the same unchallenged control mice is shown in the upper and lower panel. (**C**) Weight loss for day 5 for each group. (**D**) Analysis of oropharyngeal samples from mice at day 5 in infected groups. Ratios of E-gene to ABL1 is shown for both genomic and subgenomic RNA. Statistical comparisons are summarized as: *, p≤0.1; **, p≤0.01; **** p≤0.0001; ns, not significant. Groups displaying significant weight loss compared to uninfected mice are annotated above the points for that group.

## Discussion

Nanobodies represent a valuable alternative to monoclonal antibodies for virus neutralization^21,22^. Their ease of production, thermal stability, and robust biochemical behavior, combined with straight-forward functionalization and multimerization possibilities, make them suitable candidates for therapeutic applications.

The newly emerged SARS-CoV-2 B.1.351 variant shows nine amino acid changes in the spike protein. While biological properties of this variant remain to be determined, two amino acid substitutions in the receptor binding motif are of potential concern for vaccine efficacy, neutralization efficiency of existing monoclonal antibody therapies, and altered dynamics of virus spread^23–25^. The novel nanobody described here, Fu2, can effectively neutralize the B.1.351 variant and therefore has a strong and unique potential to form the basis for development of therapeutic interventions against this emerging variant of concern.

Fu2 is also capable of neutralizing SARS-CoV, highlighting its specificity for a more conserved epitope. However, the potency was significantly reduced for the monomeric nanobody, likely because of decreased affinity to the RBD of this virus. That the dimeric constructs significantly increased potency further supports this hypothesis. A similar observation was made for VHH-72 originally developed against SARS-CoV, but with some neutralization potential in dimeric form against SARS-CoV-2^26,27^.

Nanobodies typically have short serum half-lives, ranging from 1-3 hours^28^, compared to 6-8 days of IgG1^29^. This characteristic can dramatically influence applicability for virus neutralization. Accordingly, for a Fu2-Ty1 heterodimer, we could not detect any protective effect when administered prophylactically, while therapeutic administration significantly delayed disease onset and severity (Fig. S7). In contrast, a half-life extended Fu2-Alb1 heterodimer showed therapeutic efficacy against ‘wild-type’ and B.1.351 (Fig. 4), suggesting that nanobody-based constructs have the potential for SARS-CoV-2 antiviral therapy.

The RBD-specific nanobody Fu2 displays a particularly unique mode of virus neutralization by rapidly inducing stable spike trimer dimers that are incapable of binding to ACE2. In contrast to spike trimer dimers reported for the full-length spike that are based on S1-S1 interactions placing the trimers ‘side-by-side’^30^, Fu2 induced a dimer in a ‘head-to-head’ orientation with the C-termini at the distal ends. Targeting two epitopes increases the binding surface area, but, more importantly, when locked in such a conformation the function of each spike is sterically blocked both by the nanobody and by the other spike in such a way that the whole complex must dissociate for the spike to regain its function.

The interaction between Fu2 and the RBD is highly unusual, and for a monomer to induce antigen dimerization in this manner, two copies of an antigen must be bound at different epitopes with precisely the right spatial arrangement. This novel mode of antigen binding may be uncommon in naturally elicited antibody repertoires but could be achievable by structure-based antibody design and might present a general strategy for potent immobilization of viral glycoproteins and other antigens. It remains to be seen whether ‘designer’ bispecific monomeric nanobodies can push the potency envelope beyond what is currently attainable.

## Materials and Methods

### Cells and viruses

HEK293T cell (ATCC-CRL-3216 and ATCC CRL-11268) and Vero E6 cells (ATCC-CRL-1586) were maintained in DMEM (Gibco) supplemented with 10% fetal calf serum and 1% penicillin-streptomycin in a humidified incubator with 5% CO_2_ at 37°C. Calu-3 cells were additionally supplemented with nutrient mixture F12 (Thermo Fisher Scientific). The HEK293T cell line was further engineered to overexpress human ACE2 by lentiviral transduction^14^. All cell lines used for the experiments were negative for mycoplasma as determined by PCR. Infectious SARS-CoV-2^19^ and the B.1.351^1^ isolate used for PRNTs and the challenge experiment in Fig. S7 were propagated in Vero E6 cells and titrated by plaque assay. Viruses used for the challenge in Fig. 4 were propagated in Calu-3 cells, quantified by qPCR, and titrated by plaque assay in Vero E6 cells. Challenge stocks for experiments shown in Fig. 4 were confirmed by Sanger sequencing to harbor no high frequency cell culture adaptation mutations in spike.

### Proteins and probes

The plasmid for the expression of the SARS-CoV-2 prefusion-stabilized spike (2P) ectodomain was a kind gift from the McLellan lab^6^. The plasmid was used for transient transfection of FreeStyle 293F cells using the FreeStyle MAX reagent (Thermo Fisher Scientific). The spike trimer was purified from filtered supernatant on Streptactin XT resin (IBA Lifesciences) or Ni-NTA resin and purified by size-exclusion chromatography on a Superdex 200 (Cytiva). For cryo-EM a similar construct with 6 proline substitutions (S6-P) was used^33^, and the protein was purified on His-Pur Ni-NTA resin (Thermo Fisher Scientific) followed by size-exclusion chromatography. The RBD domain was cloned upstream of a sortase A motif (LPETG) and a 6xHIS tag. The plasmid was used for transient transfection of FreeStyle 293F cells as described above. The protein was purified from filtered supernatant on His-Pur Ni-NTA resin followed by size-exclusion chromatography on a Superdex 200. The albumin binding nanobody Alb1 was described earlier^20^ and the sequence was obtained from WO/2006/122787. Nanobodies were cloned in the pHEN plasmid with a C-terminal sortase motif (LPETG) and a 6xHIS tag. BL21 *E. coli* were transformed with this plasmid, and expression was induced with 1 mM IPTG at OD600 = 0.6 and cells were grown overnight at 30 °C. Nanobodies were retrieved from the periplasm by osmotic shock and purified on Ni-NTA resin and size-exclusion chromatography.

Sortase A 5M was produced as described before in BL21 *E. coli* and purified by Ni-NTA and size exclusion chromatography^3^. Fluorescent spike ectodomain was generated by first attaching dibenzocyclooctyine-*N*-hydroxysuccinimidyl ester (DBCO-NHS) to the spike trimer in a 3:1 molar ratio, before attaching AbberiorStar-635P-azide by click-chemistry. The final product was purified from unreacted DBCO and fluorophore on a PD-10 desalting column. The biotinylated RBD was generated using sortase A and amine-PEG3-biotin as a nucleophile. The reaction was performed with 50 µM RBD, 5 µM sortase A 5M, and 8 mM amine-PEG3-biotin in 50 mM Tris pH 7.5, 150 mM NaCl, for 6 hours at 4 °C. Sortase A and unreacted RBD was removed on Ni-NTA resin and excess nucleophile was removed by two consecutive purifications on PD-10 desalting columns.

### Alpaca immunization, library generation and nanobody isolation

One adult female alpaca (*Funny*) at PreClinics, Germany, was immunized four times in a 60-day immunization schedule. Each immunization consisted of two injections. For the first immunization, 200 µg of prefusion stabilized spike and 200 µg of S1+S2 domain (Sino Biologicals) was used. Remaining immunizations each consisted of one injection with 200 µg RBD and one injection with 200 µg prefusion stabilized spike, both produced in Freestyle 293F cells as described above. Serum from the final bleed at day 60 neutralized SARS-CoV-2 pseudoviruses with an ID50 >20 000. The animal study protocol was approved by the PreClinics animal welfare officer commissioner and registered under the registration No. 33.19-42502-05-17A210 at the Lower Saxony State Office for Consumer Protection and Food Safety—LAVES and is compliant with the Directive 2010/63/EU on animal welfare. The nanobody phage library was generated as described^14^. Phage display was performed on RBD immobilized on magnetic beads.

### Dimer generation using click chemistry

To generate homo- and hetero-dimers, the different nanobodies were first site-specifically functionalized on the C-terminus using sortase A with either an azide or a dibenzocyclooctine (DBCO) as described in detail here^3^. In brief, for functionalization with DBCO, 70 µM of nanobody was incubated with 5 µM sortase A, 8 mM DBCO-amine (Sigma-Aldrich) in 150 mM NaCl, 50 mM Tris pH 7.5 and 10 mM CaCl_2_ for 3 h at 25 °C. To functionalize the nanobody with an azide, 70 µM of nanobody, was incubated with 5 µM sortase A 5M, 10 mM 3-azido-1-propanamine (Sigma-Aldrich #762016) in 150 mM NaCl, 50 mM Tris pH 7.5 and 10 mM CaCl_2_ for 3 h at 25 °C. In both reactions, unreacted nanobody, sortase A and excess nucleophile were removed using Ni-NTA resin and PD-10 columns or size-exclusion chromatography. The click reaction was initiated by mixing azide and DBCO labeled nanobody in a 1:1 molar ratio for 48 h at 4 °C. Dimers were purified from unreacted monomeric nanobody by size-exclusion chromatography on a Superdex S200 16/600 column (Cytiva).

### Neutralization assays

Pseudotyped virus neutralization assay: Pseudotyped viruses were generated by co-transfection of HEK293T cells with plasmids encoding firefly luciferase, a lentiviral packaging plasmid (Addgene cat#8455), and a plasmid encoding either the SARS-CoV or SARS-CoV-2 spike protein, both harboring a C-terminal truncation of 18 amino acids. Media was changed 12-16 h post-transfection, and pseudotyped viruses were harvested at 48 and 72 hours, filtered through a 0.45 µm filter and stored at −80 °C until use. Pseudotyped viruses sufficient to generate 100,000 relative light units (RLU) were incubated with serial dilutions of nanobody for 60 min at 37 °C. 15,000 HEK293T-ACE2 cells were then added to each well, and the plates were incubated for 48 h at 37 °C. Luminescence was measured using Bright-Glo (Promega) on a GM-2000 luminometer (Promega) with an integration time of 0.3 s.

Plaque reduction neutralization test: One day before the experiment, 100,000 Vero E6 cells were seeded per well of a 24-well plate. A 3-fold serial dilution of the nanobody constructs starting at 10 µg/ml were incubated with 100 plaque-forming units (PFU) of the first Swedish clinical isolate of SARS-CoV-2 or variant B.1.351 for 1 h at 37°C. The nanobody-virus mixture was then added to the cells and incubated at 37 °C for 1 h. Cells were washed with PBS, and an overlay of 1% CMC/DMEM supplemented with 2% FBS was pipetted into the wells. Three days post-infection, cells were fixed in 10% formaldehyde, followed by staining with crystal violet. Plaques were counted and inhibition determined by comparing to control wells on the same plate.

### Flow cytometry

HEK293T-hACE2 cells were trypsinized and fixed in 4% formaldehyde in PBS for 20 min. Cells were stained with spike-AbberiorStar-635P not premixed or premixed with Fu2 or control nanobody. Fluorescence was quantified using a BD FACSCelesta and the FlowJo software package.

### Surface plasmon resonance

Binding kinetics were determined by surface plasmon resonance using a Biacore 2000. All experiments were performed at 25 °C in a running buffer of 10 mM HEPES, 150 mM NaCl, pH 7.4, and 0.005% Tween-20 (v/v). Site-specifically biotinylated RBD was immobilized on streptavidin sensor chips (Series S sensor Chip SA, GE Healthcare) to a level of ~200 resonance units (RU). A 2-fold dilution series of the nanobodies was injected at a flow rate of 30 µl/min (association 180 s, dissociation 900 s), and the immobilized RBD was regenerated using 0.1 M glycine-HCL buffer pH 2 for 2x 10 seconds. Data were analyzed using BIAevaluation Software and fitted using the 1:1 Langmuir model with mass transfer.

### Next generation sequencing

Next generation sequencing on pre- and post-enrichment nanobody libraries was performed as described in Hanke et al. 2020^14^ on an Illumina MiSeq.

### Gel-filtration shift assays

For the gel-filtration shift assay, 80 µg of prefusion stabilized spike protein was incubated with or without nanobody for 15 minutes and was run on a Superose 6 increase 10/300 GL (Cytiva), and absorbance at 280 nm was detected. Sample and injection volumes were kept identical between runs to obtain comparable elution profiles.

### Fluorescent pseudotyped lentivirus brightness measurements

To generate fluorescent lentiviral particles enveloped with the spike protein of the SARS-CoV-2 coronavirus (PSV-GFP), HEK 293T cells were reverse-transfected with 3 plasmids: psPAX2 (a gift from Didier Trono, Addgene #12260), pEGFP-Vpr (obtained through the NIH HIV Reagent Program, Division of AIDS, NIAID, NIH: pEGFP-Vpr, ARP-11386, contributed by Dr. Warner C. Greene) and pCMV14-3X-Flag-SARS-CoV-2 S (a gift from Zhaohui Qian, Addgene #145780) at 1:0.5:2 molar ratio using Lipofectamine™ 2000 (0.6 ul per 1 ug DNA; Invitrogen™, 11668019); growth medium was refreshed 16 h after transfection. Virus-rich supernatant was collected twice over the next 48 h and, if needed, stored at 4 °C prior to the concentration. Supernatants were pooled, mixed with the Lenti-X™ Concentrator (Takara Bio, 631231) at 3:1 ratio (v:v) and incubated overnight at 4 °C. The solution was then spun down at 1’500 x g for 45 min at RT, the pellet was resuspended in sterile 1x PBS (Sigma-Aldrich, D8537) and stored at −80 °C. PSV-GFP was incubated with 100 µg/ml Ty1, Fu2 or PBS as control for 1 h. The mixtures were transferred to an 8-well glass bottom Ibidi chamber. Fluorescence Correlation Spectroscopy (FCS) measurements were performed using Zeiss LSM 780 microscope. 488 nm argon ion laser was used to excite GFP. 40x/1.2 NA water immersion objective was used to focus the light in solution. 40-50 curves were taken (5 s each) in solution where PSV-GFP particles diffuse freely. Laser power was set to 0.1% of the total laser power that corresponds to ≈2 μW. Curves were then fitted with FoCuS-point software^34^ to extract number of diffusing species and brightness of single diffusing species.

### Cryo-EM sample preparation and imaging

Spike trimer (1.8mg/ml) S6-P and Fu2 (2.1mg/ml) were mixed in a 1:6 molar ratio followed by incubation on ice for 10 minutes. Prior to cryo-EM grid preparation, grids were glow-discharged with 25 mA for 2 minutes using an EMS 100X (Electron Microscopy Sciences) glow-discharge unit. CryoMatrix® holey grids with amorphous alloy film (R 2/1 geometry; Zhenjiang Lehua Technology Co., Ltd) were used. 3 μl aliquots of sample solutions were applied to the grids and the grids with sample were then vitrified in a Vitrobot Mk IV (Thermo Fisher Scientific) at 4°C and 100% humidity (blot 10 s, blot force 3, 595 filter papers (Ted Pella Inc.)). Cryo-EM data collection was performed with EPU (Thermo Fisher Scientific) using a Krios G3i transmission-electron microscope (Thermo Fisher Scientific) operated at 300 kV in the Karolinska Institutet’s 3D-EM facility (https://ki.se/cmb/3d-em). Images were acquired in 165 kx nanoprobe EFTEM SA mode with a slit width of 10 eV using a K3 Bioquantum. Exposure time was 1.5 s during which 60 movie frames were collected with a fluency of 0.81 e-/Å2 per frame (Table S1). Motion correction, CTF-estimation, Fourier binning (to 1.01 Å/px), picking and extraction in 600-pixel boxes (size threshold 400 Å, distance threshold 100 Å, using the pretrained BoxNet2Mask_20180918 model) were performed on the fly using Warp^35^. Due to preferential orientation, we collected a part of the dataset with a 25-degree stage tilt.

A total of 14,081 micrographs were selected based on an estimated resolution cut-off of 4 Å and defocus below 2 microns as estimated by Warp. Furthermore, Warp picked 1,035,962 particles (dimer of trimeric spike with Fu2) based on above-mentioned criteria. Extracted particles were imported into cryoSPARC v3.1^36^ for 2D classification, 3D classification and non-uniform 3D refinement. After 2D classification, clean classes with high-resolution features (and with characteristic trimeric spike views) were retained and used to generate ab-initio 3D reconstructions. In total 277,372 particles were retained for refinement steps and these particles were processed with C1 symmetry (image processing scheme Fig. S5). These particles were further processed using heterogeneous refinement that resulted in a reconstruction with high-resolution structural features in the core of the spike. One round of homogeneous refinement was followed by non-uniform refinement. All final reconstructions were analysed using 3D-FSC ^37^ and there were moderate anisotropy in the full map reconstructions while the localized reconstruction displayed no significant anisotropy (Fig. S6A and S6B). All CTF refinements were per-particle CTF refinements interspersed with global aberration correction (beamtilt, trefoil, tetrafoil and spherical aberration). Please see Table S1 for data collection and processing statistics and the respective cryo-EM data processing schemes.

For the Spike-Fu2 dimer interface we used a particle set with partial-signal subtraction of all parts except for Fu2-RBD dimer interface (containing two RBDs and two Fu2). From this we performed local reconstruction (non-uniform). The local reconstruction resulted in a map with 2.89Å overall resolution as compared to 3.2Å overall resolution for the full map.

### Cryo-EM model building and structure refinement

The structure of the spike protein trimer PDB: 7KSG^5^ was used as a starting model for model building. A homology model for Fu2 was generated by SWISS-MODEL^38^ with PDB: 5LHN as template (chain B)^39^. Structure refinement and manual model building were performed using Coot^40^ and PHENIX ^41^ in interspersed cycles with secondary structure, Ramachandran, rotamers and bond geometry restrains. Structure figures and EM density-map figures were generated with UCSF ChimeraX^42^ and COOT, respectively. Please see Table S2 for refinement and validation statistics.

### SARS-CoV-2 challenge experiments

K18-hACE2 transgenic mice were purchased from Jackson laboratories and maintained as a hemizygous line. Experiments were conducted in BSL3 facilities at the Comparative Medicine department (KM-F) at Karolinska Institutet. Ethics for studies of virus infection and therapeutic intervention were obtained from the Swedish Board of Agriculture (10513-2020). Mice were administered nanobodies as described in the main text and challenged intranasally with 86 PFU SARS-CoV-2 (‘wild-type’, Swedish isolate) or 100 PFU B.1.351 in 40 µl PBS following isoflurane sedation. The challenge in Fig. S7 was performed with 1000 PFU of SARS-CoV-2 (Swedish isolate) propagated in Vero E6 cells. Oropharyngeal sampling was performed at the indicated timepoints under light anesthesia with isoflurane. Weight and general body condition were monitored daily until weight drop started, whereupon mice were monitored twice daily. During the experiment, weight loss, changes in general health, breathing, body movement and posture, piloerection and eye health were monitored. Mice were typically sacrificed when they achieved 20% weight loss. However, some mice were sacrificed before losing 20% in body weight, when movement was greatly impaired and/or they experienced difficulty breathing that was considered to reach a severity level of 0.5 on Karolinska Institutet’s veterinary plan for monitoring animal health. The weight loss in response to infection was highly reproducible. In Fig. 4 data from 50% of the challenged and untreated animals are historical controls from previous experiments performed under identical conditions.

### RNA Extraction and RT-qPCR for SARS-CoV-2 detection

Viral RNA was isolated from buccal swabs collected on dpi 6 and stored in 500 µl of TRIzol™ Reagent (Invitrogen). Total RNA extractions from buccal swab samples were performed using an adapted TRIzol™ manufacturers protocol with a 45 min precipitation step at −20° C. RNA pellets were resuspended in 20 µl of RNase-free water.

RT-PCR reactions were performed using 4 µl of resuspended RNA in a 20 µl reaction volume using the Superscript III one step RT-qPCR system with Platinum Taq Polymerase (Invitrogen) with 400 nM concentrations of each primer and 200 nM of probe. Primers and probes for the CoV-E gene target were as previously described ^43^. Primers and probes for the ABL1 target were adapted from Ishige et al.^44^ to enable detection of the murine homolog: (ABL1_ENF1003_deg: 5’-TGGAGATAACACTCTCAGCATKACTAAAGGT-3’, ABL1_ENR1063: 5’-GATGTAGTTGCTTGGGACCCA-3’, ABL1_ENPr1043_deg: 5’-HEX-CCATTTTTSGTTTGGGCTTCACACCATT-BHQ1-3’). The CoV-E and ABL1 TaqMan assays were run in multiplex. Detection of the subgenomic CoV-E target was adapted from Wölfel et al.^45^, using a leader/E gene junction specific forward primer: (sgEjunc_SARSCoV2_F: 5’-CGATCTCTTGTAGATCTGTTCTCTAAACG-3’). All oligonucleotides were synthesized by Eurofins Genomics.

Thermal cycling conditions for all assays consisted of RT at 55 °C for 10 min, denaturation at 95 °C for 3 min, and 45 cycles of 95C, 15s and 58C, 30s. Reactions were carried out using a CFX96 Connect Real-Time PCR Detection System (Bio-Rad) following manufacturer instructions. To generate standard curves, a synthetic DNA template gBlock (Integrated DNA Technologies) was transcribed using the mMessage mMachine™ T7 Transcription Kit (Invitrogen) and serially diluted. To reduce sampling-related variability, SARS-CoV-2 RNA copies were normalized by ABL1 copies, and this ratio was used for comparisons. ABL1 copies were not significantly different between groups.

## Acknowledgments

Experiments with replication-competent SARS-CoV-2 were performed in the Biomedicum BSL3 core facility, Karolinska Institutet. We thank Jonas Klingstrom for providing Calu-3 cells and sharing the Swedish SARS-CoV-2 isolate, and Alex Sigal from the Africa Health Research Institute for providing the B.1.351/501Y.V2 isolate. All cryo-EM data were collected in the Karolinska Institutet’s 3D-EM facility. We thank Agustin Ure for assistance with figure generation and Tomas Nyman (Protein Science Facility at KI) for providing access to SPR instruments.

## Funding

LH was supported by the David och Astrid Hageléns stiftelse, the Clas Groschinskys Minnesfond and a Jonas Söderquist’s scholarship. This project has received funding from the European Union’s Horizon 2020 research and innovation program under grant agreement No. 101003653 (CoroNAb), to BM and GMM. BMH is supported by the Knut and Alice Wallenberg Foundation (KAW 2017.0080 and KAW 2018.0080). The work was supported by project grants from the Swedish Research Council to BMH (2017-6702 and 2018-3808), BM (2018-02381) and to GMM (2018-03914 and 2018-03843).

## Author contributions

Conceptualization: LH, BM, GMM. Methodology: LH, DJS, JMC, BMH, BM, GMM. Investigation: LH, HD, DJS, LPV, EU, AMM, VK, BP, ES, NS, AP, CK, GKP, BMH. Data Curation: LH, BM. Visualization: LH, HD, DJS, BMH, BM. Funding acquisition: LH, BMH, BM, GMM. Supervision: LH, DJS, OFC, JMC, BMH, BM, GMM. Writing – original draft: LH, HD. Writing – review & editing: LH, HD, DJS, LPV, EU, AMM, VK, ES, GKP, JMC, BMH, BM, GMM.

## Competing interests

LH, DJS, LPV, BM, and GMM are listed as inventors on a patent application describing SARS-CoV-2 nanobodies.

## Data and materials availability

The cryo-EM density maps have been deposited in the Electron Microscopy Data Bank under accession codes EMD-12561 (dimer of spike trimer + 6 Fu2) and EMD-12465 (localized reconstruction of 2 RBDs + 2 Fu2). The atomic coordinates have been deposited in the Protein Data Bank under IDs 7NS6 (dimers of spike trimers + 6 Fu2) and 7NLL (localized reconstruction of 2 RBDs + 2 Fu2). All nanobody sequences will be made available on public databases upon publication.

## Supplementary figures and tables

**Fig. S1.**
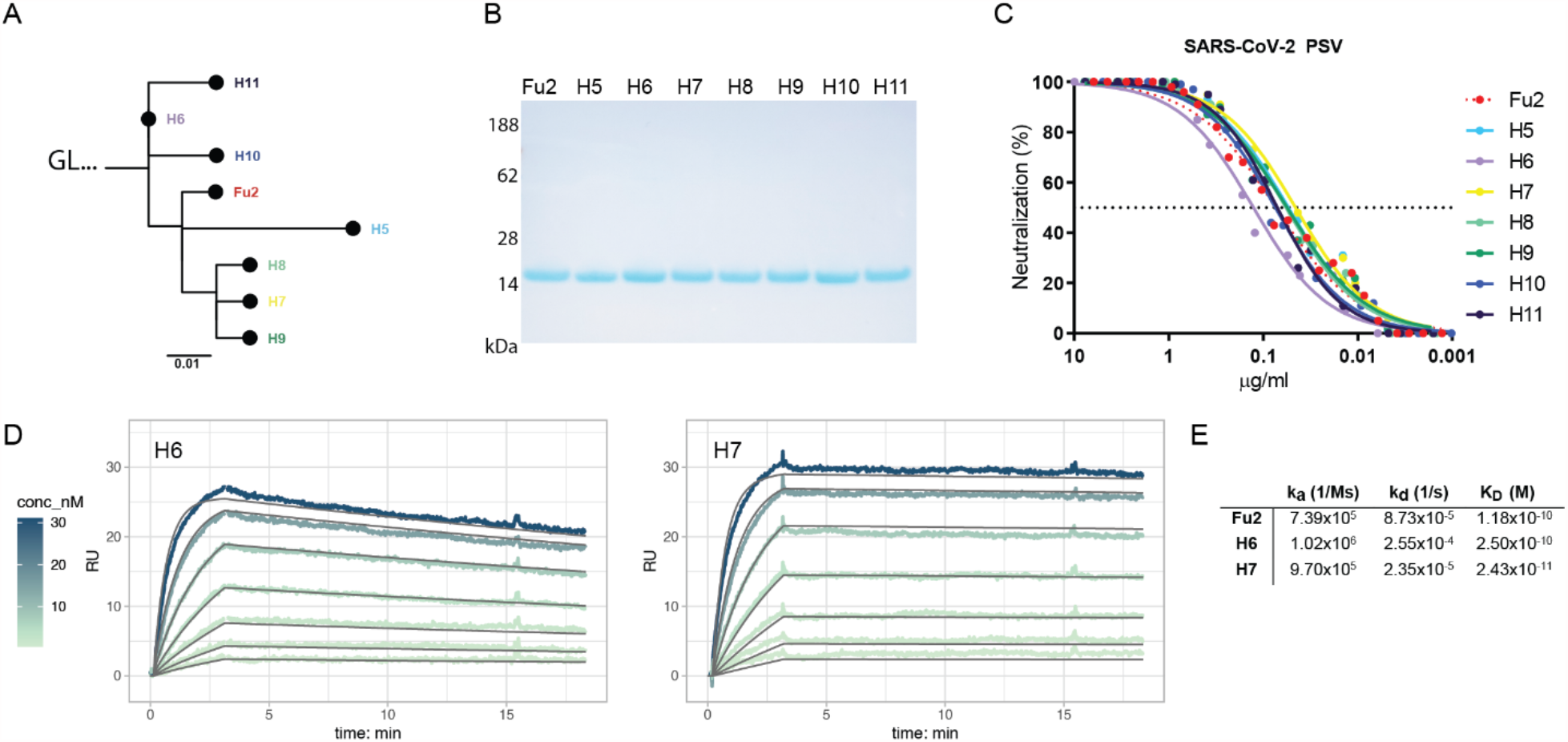
Fu2 variants identified by next-generation sequencing (NGS) display similar neutralization potential and binding kinetics. (**A**) Average difference of Fu2 variants is shown as a tree diagram. (**B**) Fu2 and seven Fu2-variants were expressed in *E. coli*, purified, and analyzed by SDS-PAGE and Coomassie staining. (**C**) A dilution series of Fu2 and its variants were incubated with SARS-CoV-2 pseudotyped lentivirus (PSV) for 1 hour at 37 °C before infecting HEK293T-hACE2 cells. Neutralization (in %) compared to the untreated PSV is shown. (**D**) Binding kinetics of Fu2 and variants to the RBD were measured by surface plasmon resonance (SPR). Site-specifically biotinylated RBD was immobilized on streptavidin sensor chips, and kinetics for a dilution series of the indicated monomeric nanobodies were measured. Sensorgrams are color-coded based on concentration. The fit is based on the 1:1 Langmuir model and is shown in dark grey solid lines. (**E**) Kinetic parameters of Fu2 and Fu2 variants binding to the RBD.

**Fig. S2.**
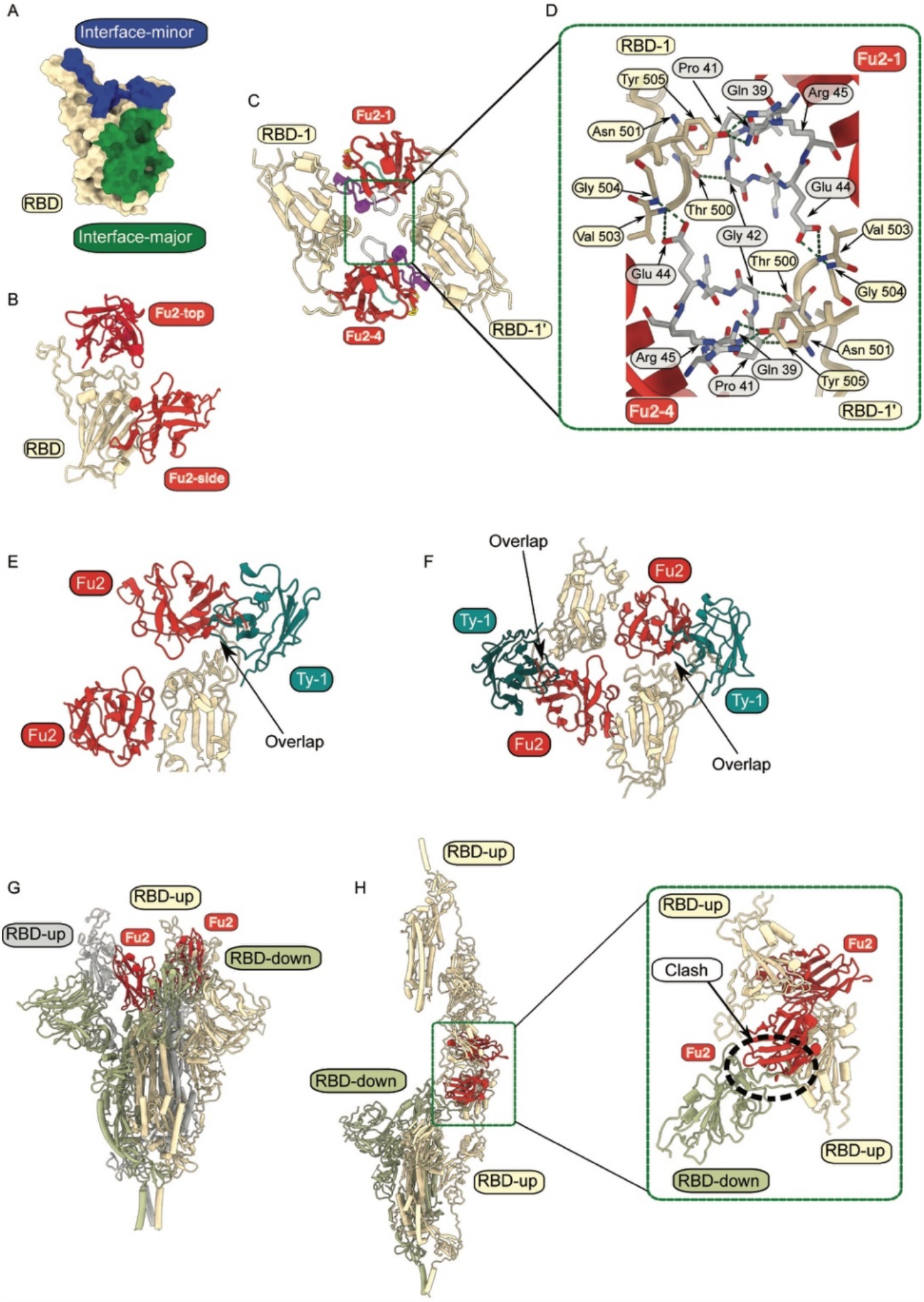
(**A**) Surface view of the RBD with interface-major and interface-minor color coded. (**B**) Relative positions of two Fu2 molecules binding the RBD. (**C)** The interaction between two RBDs and two Fu2s showing in grey the β-hairpins in the Fu2 framework region (39-45) that help to mediate the dimer-of-trimers Fu2-spike complex. (**D**) Close-up view of (C) (**E** and **F**) Assessment of Fu2-Ty1 heterodimer binding. Alignment of the Spike-Fu2 structure to the Spike-Ty1 structure (RBD in ‘up’ conformation) (PDB:6ZXN^14^). Arrows indicate partial overlap of binding surface on RBD. (**G**) Assessment of Fu2-RBD interaction in RBD-down conformation. RBD-Fu2 structure superimposed to RBD-down of spike in 2-up conformation (PDB:7A29) (**H**) Assessment of Fu2-RBD interaction in RBD-down conformation in dimeric spike. 2-up spike (PDB:7A29^13^) is shown superimposed with the spike dimer-of-trimers.

**Fig. S3.**
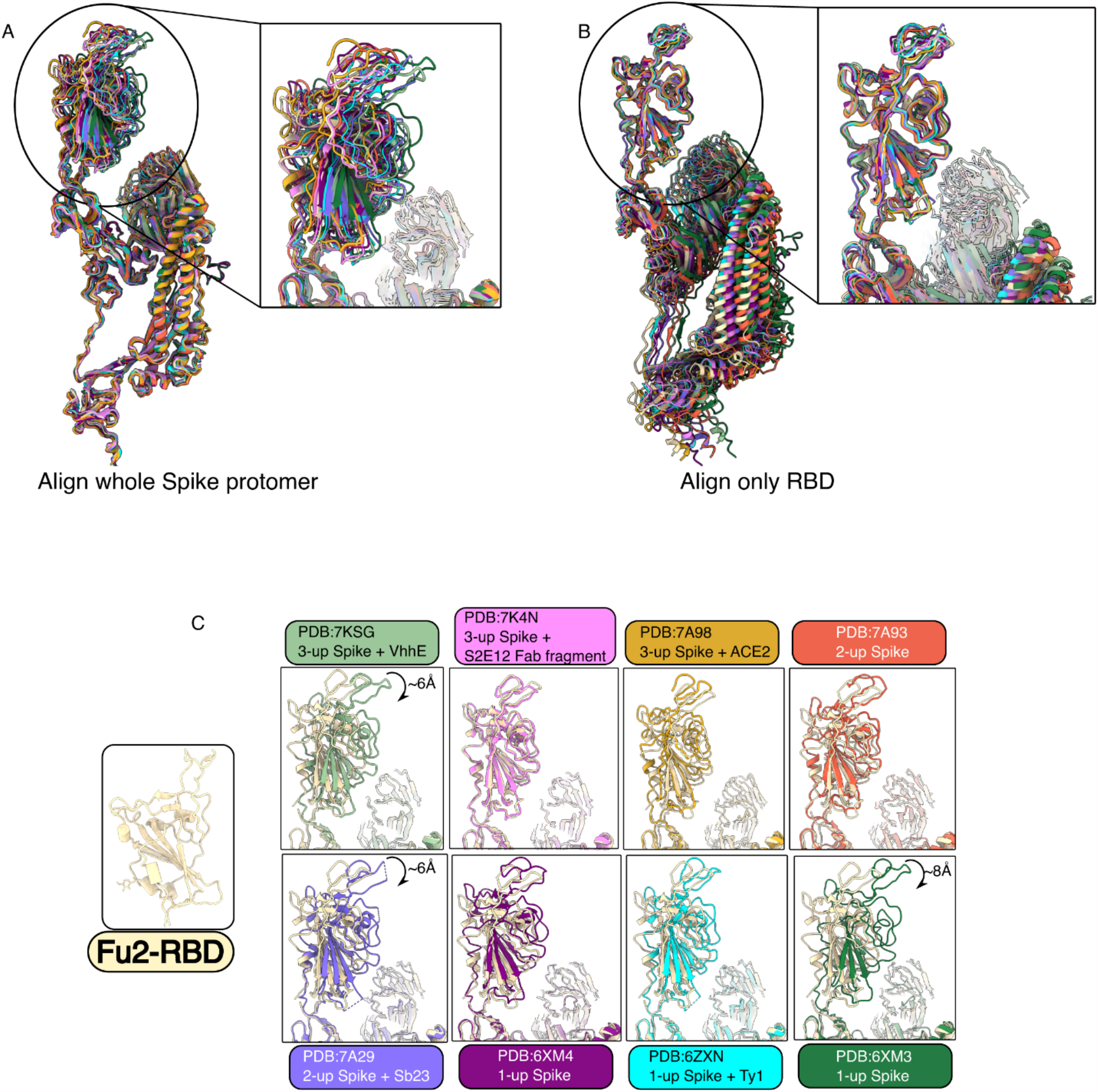
Assessment of Fu2 mediated structural changes in spike structure. (**A**) Alignment of dimeric spike trimer with spikes in 1-up, 2-up and 3-up conformation (one protomer in ‘up’ state shown). (**B**) Structural alignment of dimeric spike trimer with RBDs of 1-up, 2-up and 3-up spikes (protomer in ‘up’ conformation). (**C**) Structural comparison of RBD (Fu2-bound dimeric spike) with eight different RBDs structures (7KSG^5^, 7K4N^46^, 7A98^47^, 7A93^47^, 7A29^13^, 6XM3^48^, 6XM4^48^, 6ZXN^14^).

**Fig. S4.**
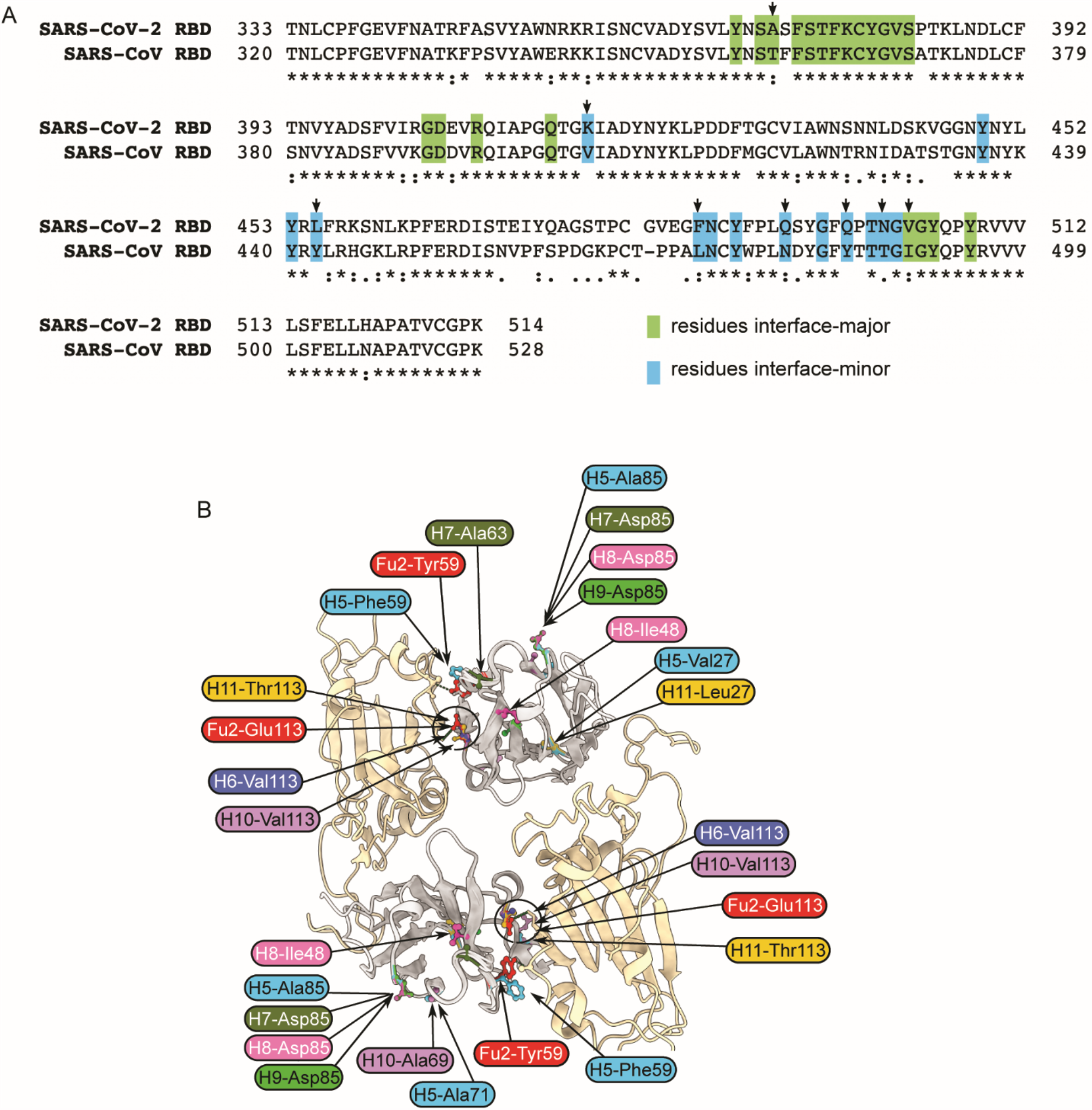
(**A**) Sequence alignment of RBDs from SARS-CoV and SARS-CoV-2. Interface-major and interface-minor residues are highlighted. Arrows indicate the different amino acids of the interface. (**B**) Fu2 variants and analysis of interface residues. Distinct residues of different variants shown as sticks (color coded).

**Fig. S5.**
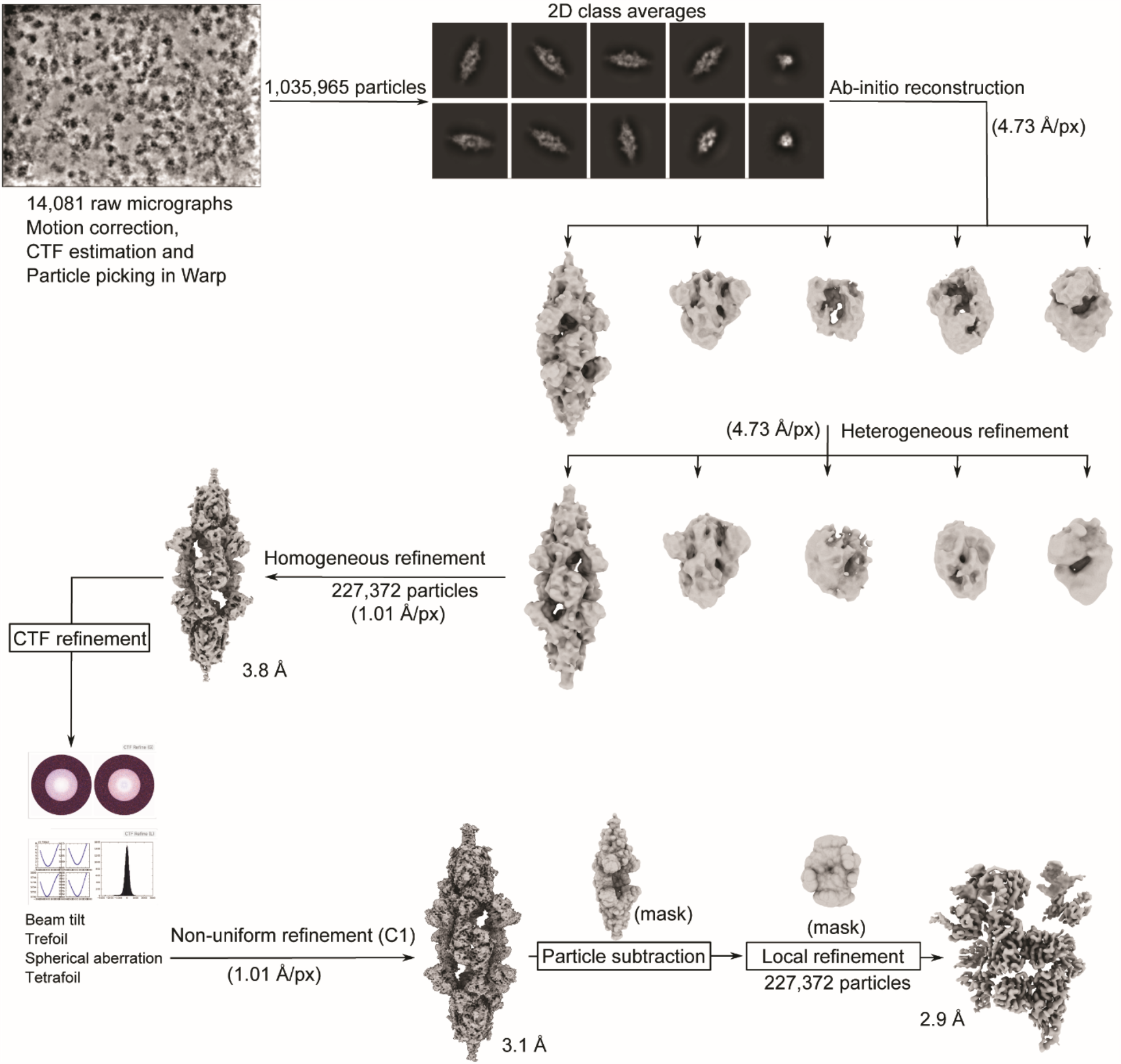
Cryo-EM image processing scheme. Particles picked by Warp^35^ were processed in cryoSPARC v3.1^36^. Representative 2D class averages are shown. Particles contributing to the clean classes were used to generate ab-initio reconstructions (five classes) followed by heterogeneous refinement. One class showing high-resolution features was refined further (C1 symmetry). For localized reconstruction, particle subtraction followed by local refinement (Non-uniform) was performed.

**Fig. S6.**
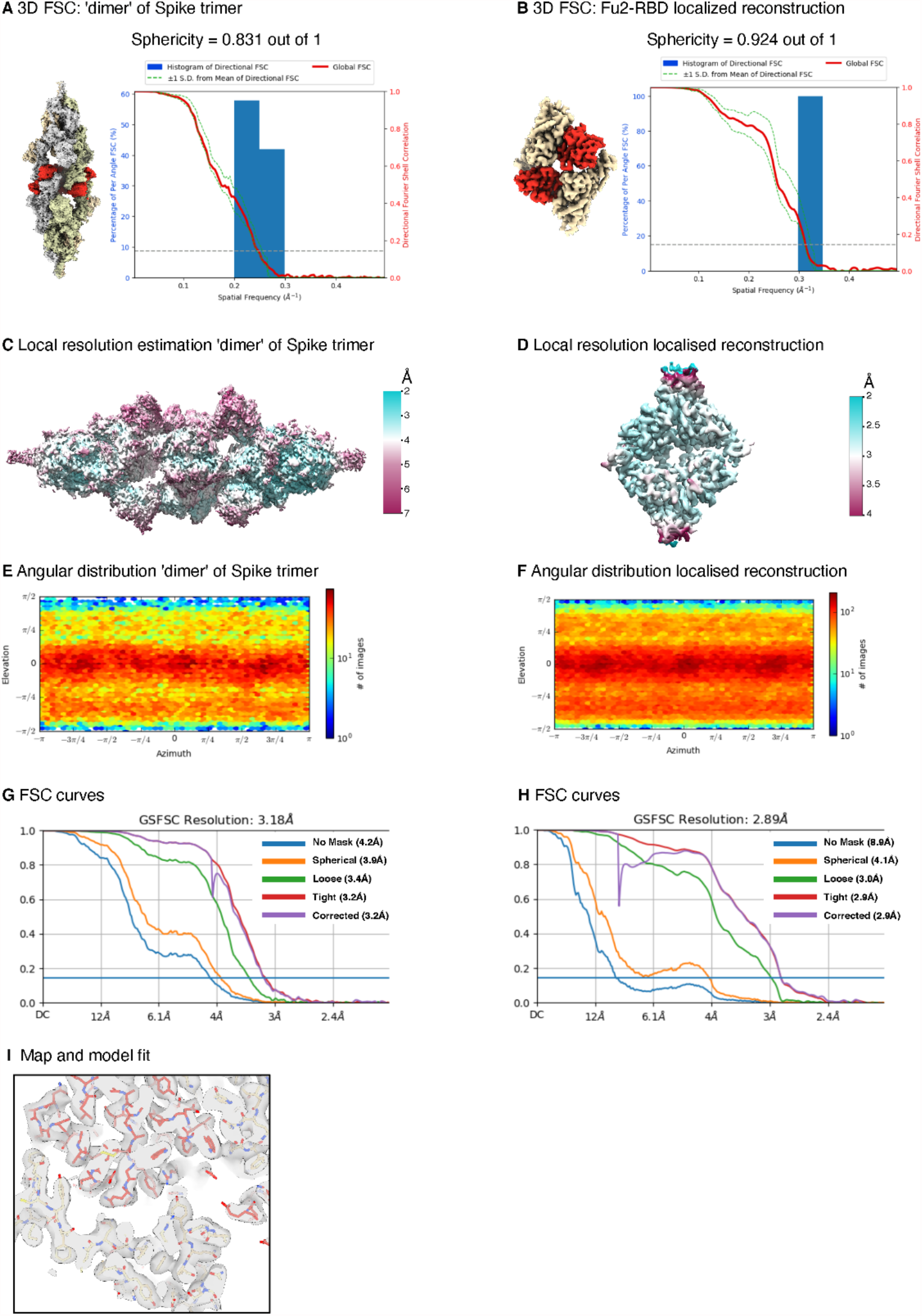
Cryo-EM validation. (**A**) and (**B**) 3D FSC^37^ and sphericity of ‘dimer’ of spike trimer and of localized map, (**C**) and (**D**) Local resolution estimation of ‘dimer’ of spike trimer and local resolution estimation of localized map (**E**) and (**F**) Angular distribution of ‘dimer’ of spike trimer reconstruction and localized reconstruction map (**G**) and (**H**) FSC curves of ‘dimer’ of spike trimer and of localized map (**I**) model-map fitting atomic resolution.

**Fig. S7.**
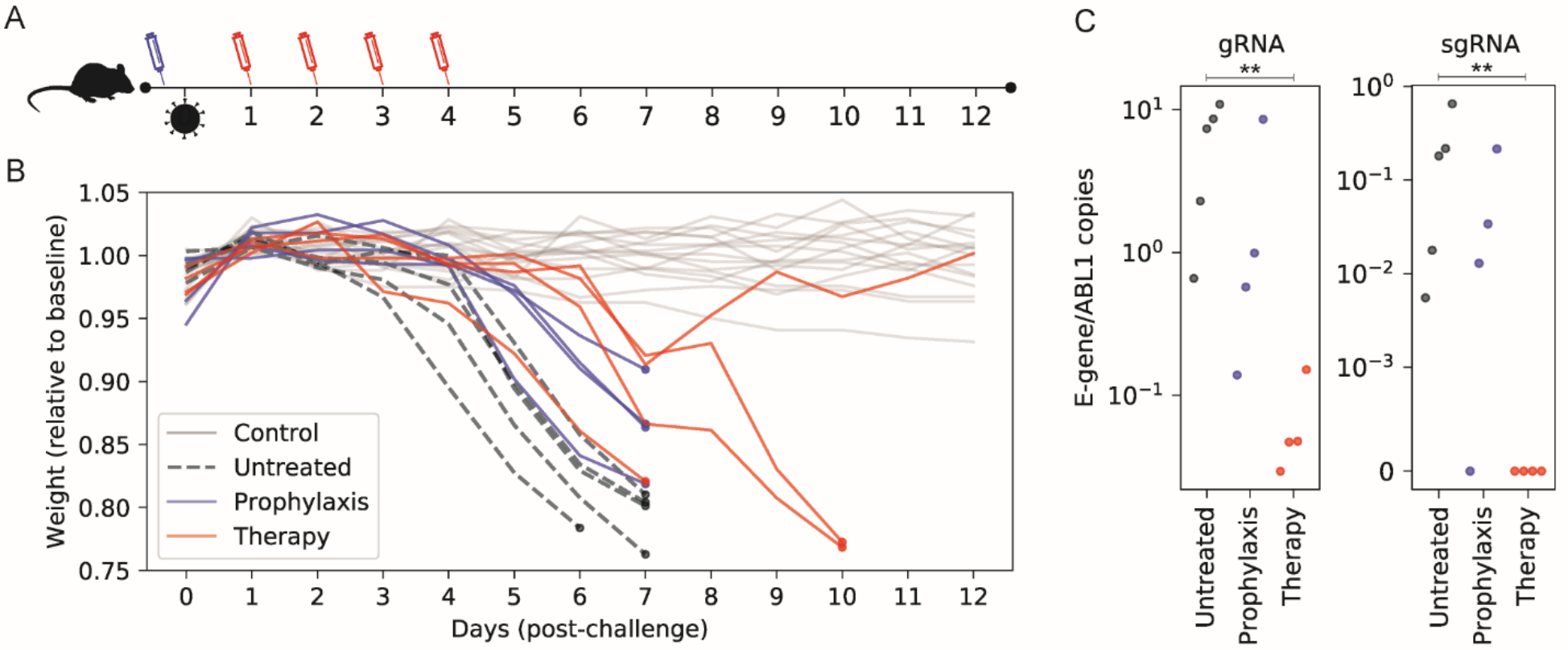
Non-half-life extended nanobody heterodimer reduces disease severity in a SARS-CoV-2 challenge model. (**A**) Timeline of the challenge experiment. K18-hACE2 transgenic mice were challenged with 1000 plaque forming units (PFU) of SARS-CoV-2 (Swedish isolate) and received prophylactic (blue) or therapeutic (red) Fu2-Ty1 at the indicated time points. (**B**) Weight of mice during the challenge experiment. The mean weight of each mouse of day 0 to day 2 served as baseline and the weight loss relative this baseline is shown. Uninfected mice are shown in grey, untreated infected mice in black, prophylactic treatment group in blue and therapeutic group in red. (**C**) Analysis of oropharyngeal samples from mice at day 6 in infected groups. Ratios of E-gene to ABL1 is shown for both genomic and subgenomic RNA (** p < 0.01, Mann–Whitney U test, one-tailed).

**Table S1.**
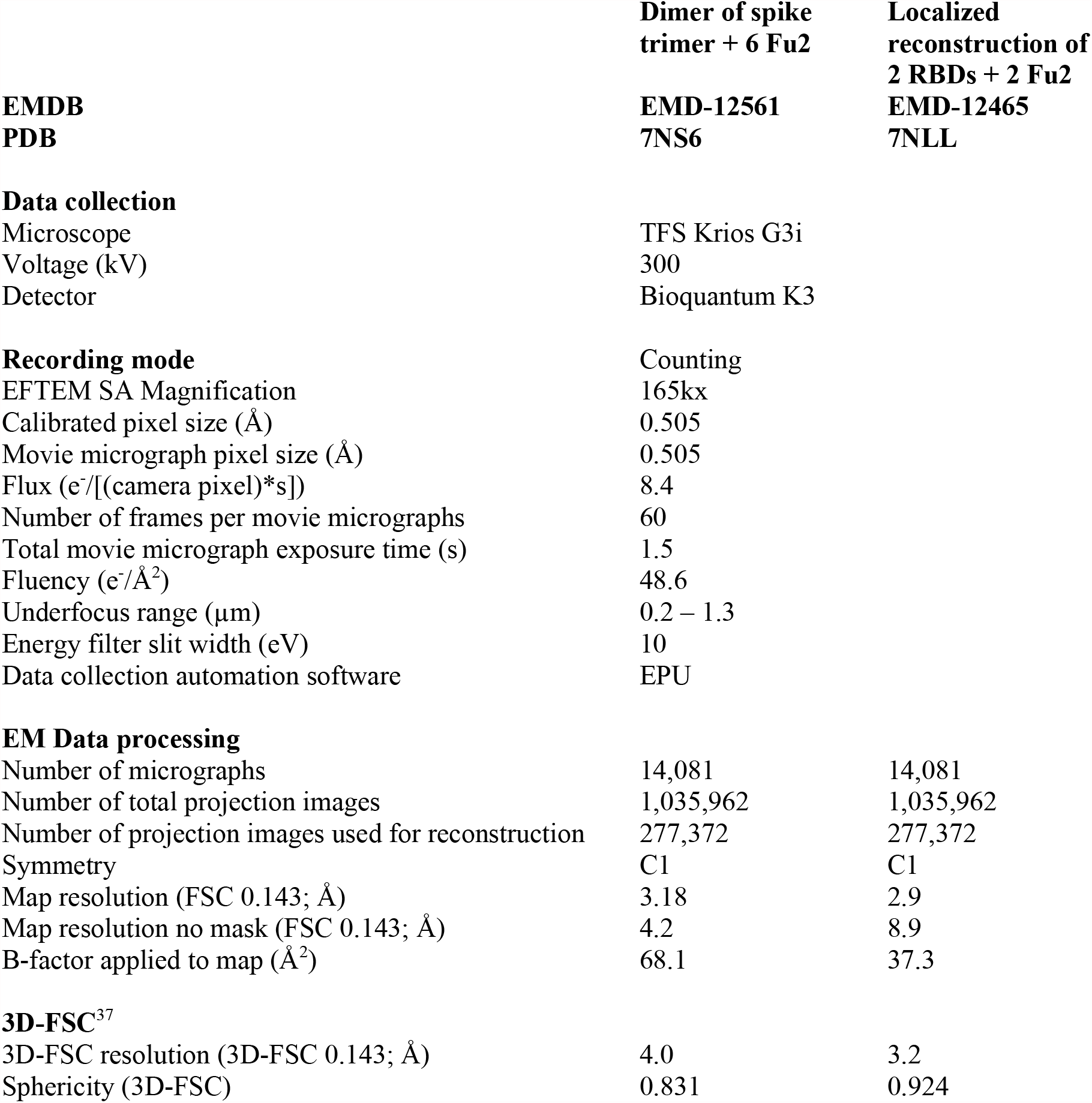
Cryo-EM data collection and processing.

**Table S2.**
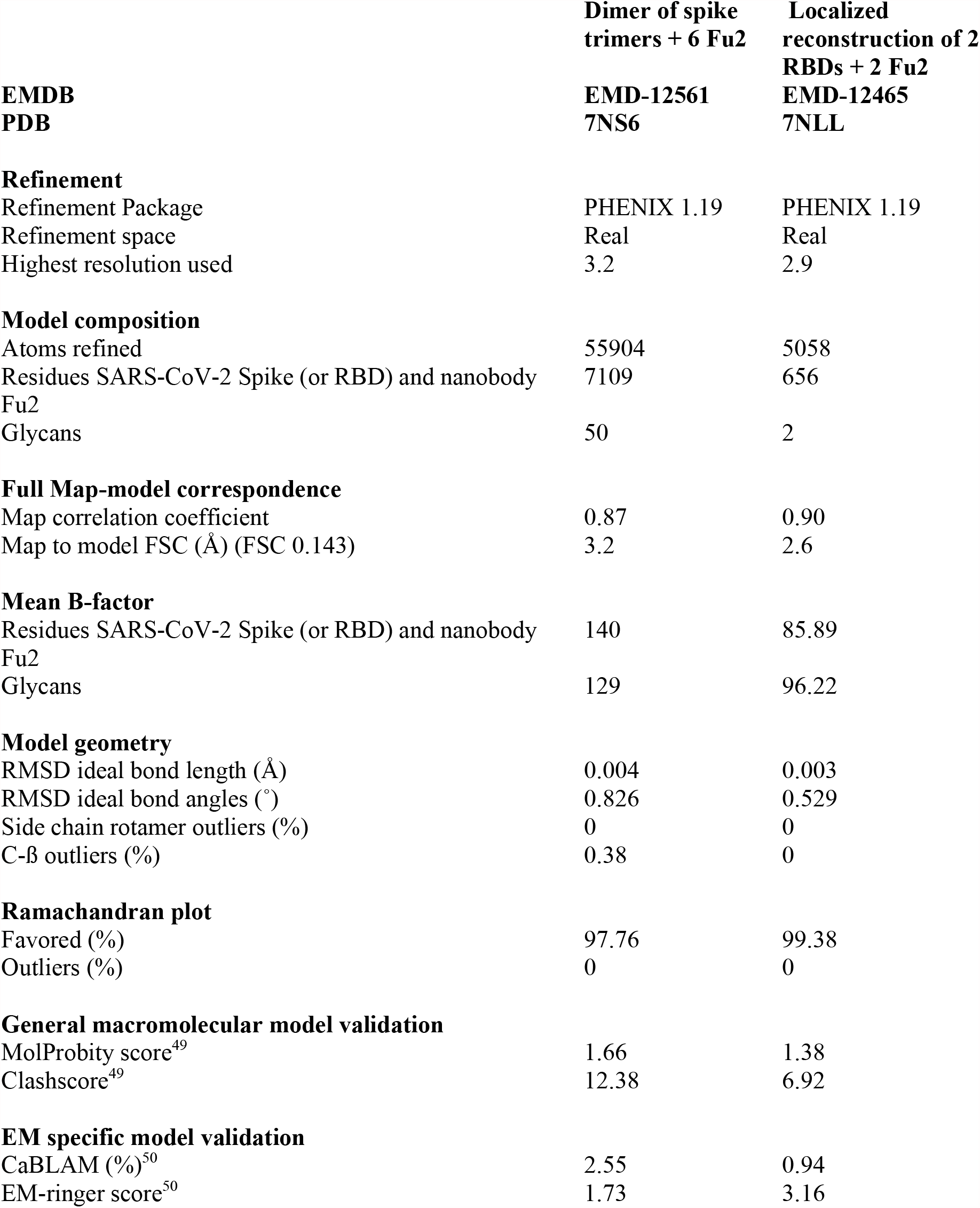
Cryo-EM model refinement and validation statistics.

| **EMDB**<br>**PDB** | **Dimer of spike**<br>**trimers + 6 Fu2**<br>**EMD-12561**<br>**7NS6** | **Localized**<br>**reconstruction of 2**<br>**RBDs + 2 Fu2**<br>**EMD-12465**<br>**7NLL** |
| --- | --- | --- |
| **Refinement** |  |  |
| Refinement Package | PHENIX 1.19 | PHENIX 1.19 |
| Refinement space | Real | Real |
| Highest resolution used | 3.2 | 2.9 |
| **Model composition** |  |  |
| Atoms refined | 55904 | 5058 |
| Residues SARS-CoV-2 Spike (or RBD) and nanobody<br>Fu2 | 7109 | 656 |
| Glycans | 50 | 2 |
| **Full Map-model correspondence** |  |  |
| Map correlation coefficient | 0.87 | 0.90 |
| Map to model FSC (Å) (FSC 0.143) | 3.2 | 2.6 |
| **Mean B-factor** |  |  |
| Residues SARS-CoV-2 Spike (or RBD) and nanobody<br>Fu2 | 140 | 85.89 |
| Glycans | 129 | 96.22 |
| **Model geometry** |  |  |
| RMSD ideal bond length (Å) | 0.004 | 0.003 |
| RMSD ideal bond angles (°) | 0.826 | 0.529 |
| Side chain rotamer outliers (%) | 0 | 0 |
| C-β outliers (%) | 0.38 | 0 |
| **Ramachandran plot** |  |  |
| Favored (%) | 97.76 | 99.38 |
| Outliers (%) | 0 | 0 |
| **General macromolecular model validation** |  |  |
| MolProbity score<sup>49</sup> | 1.66 | 1.38 |
| Clashscore<sup>49</sup> | 12.38 | 6.92 |
| **EM specific model validation** |  |  |
| CaBLAM (%)<sup>50</sup> | 2.55 | 0.94 |
| EM-ringer score<sup>50</sup> | 1.73 | 3.16 |

## Notes

### Summary of Updates

We include data showing that the Fu2 nanobody is highly protective in a mouse model of SARS-CoV-2 infection using the Swedish isolate and also the B.1.351 variant

